# Peruvian Population Genomics: Unraveling the Genetic Landscape and Admixture Dynamics of Urban Populations

**DOI:** 10.1101/2025.04.10.648256

**Authors:** Victor Borda, Omar Caceres, Cesar Sanchez, Carlos Padilla, Diego Veliz-Otani, Marla Mendes, Carolina Silva-Carvalho, Isabela Alvim, Bradley A. Maron, Mateus H. Gouveia, Peruvian Genome Project Consortium, Eduardo Tarazona-Santos, Heinner Guio, Timothy D. O’Connor

**Author notes:** Corresponding authors: Timothy D. O’Connor, Heinner Guio, and Omar Caceres.

## Abstract

Latin American populations exhibit high genetic and phenotypic diversity shaped by complex admixture histories, yet remain underrepresented in genomic research. Here, we analyze genome-wide data from 432 urban individuals across 13 regions of Peru, including 346 newly genotyped from the Peruvian Genome Project. We revealed fine-scale population structure and demographic patterns shaped by both ancient and recent events. Indigenous American ancestries in urban individuals trace back to ancient north-south interactions consisted with archaeological records, while admixture events occurring within the last 8–10 generations involved sources already admixed between distinct ancestral lineages. Identity-by-descent analyses reveal sustained gene flow in southern Peru, while effective population size trends highlight demographic stability in Lima over the past 25 generations. Sex-biased admixture patterns suggest Indigenous ancestry contribution preferentially mediated by females. These findings offer a comprehensive view of Peru’s genetic heritage, advancing our understanding of human genetic diversity and historical demographic processes in Latin America.

## Introduction

Less than 0.4% of individuals in genome-wide association studies represent Latin American populations^1^. This underrepresentation is particularly concerning given the rich cultural and genetic diversity within these populations, which have been shaped by extensive continental admixture over the past 20 generations ^2–7^ and high population structure derived from their indigenous sources ^8–10^. In particular, Peru is a case of remarkable demographic and genetic complexity. Home to approximately 33 million inhabitants and spanning 1.28 million square kilometers—an area comparable to the combined landmasses of the United Kingdom, France, and the Iberian Peninsula—Peru is one of the ancient cradles of civilization in America. With a largely Indigenous population, especially in the highlands of the Andes, it remains one of the few Western South American countries where Indigenous peoples prevail. Peruvians exhibit considerable Indigenous American genetic ancestry ^11,12^, with the average individual sharing about 50% of their genome with Indigenous groups ^8,12,13^. Nevertheless, the current evolutionary dynamics driving Indigenous ancestries in urban settings, alongside the impact of internal migrations, remain largely unexplored.

Unraveling the genetic history of Peruvians requires understanding their Indigenous American roots, which are deeply influenced by the country’s unique geography. Genetic, archaeological, and linguistic evidence suggested that Indigenous Peruvian groups were organized in a latitudinal pattern, with two interaction spheres representing the Northern and Southern mirroring Andean geography ^14,15^. Despite the profound demographic changes brought about by colonialism, present-day Indigenous American groups still exhibit genetic signatures of this North-South pattern ^9,16^. However, it remains unknown whether individuals living in urban areas follow the same pattern.

The European invasion and the arrival of enslaved Africans in the 1530s profoundly influenced recent Peruvian history. It transformed its cultural landscape and triggered significant socioeconomic changes (e.g., mines and agriculture), resulting in internal migration events ^17^. Following nearly three centuries as a prominent Spanish Viceroyalty, Peru gained independence in 1821, giving rise to a supposedly independent Republic ^18^. During the colonial period, tens of thousands of enslaved Africans were forcibly transported across the Atlantic to Panama and finally shipped along the Pacific coast in Peru to labor in mines and plantations. The Pacific Coast regions supported major economic activities related to sugar cane, cotton, and rice plantations, predominantly reliant on enslaved African labor^19^, later replaced by East Asian (Chinese and Japanese) workers after the abolition of slavery in 1854^20,21^. Thus, the last 500 years of Peruvian history have been part of a process in American social history commonly known as *mestizaje*, a term scrutinized today from different perspectives, including population genetics ^22^.

In the Amazonian region, during the late 19th and 20th centuries, the rubber industry left profound negative effects on local indigenous communities and favored internal immigration from other Peruvian regions to Amazonian cities ^23^. In addition, continual rural-to-urban migration, a trend prevalent in many Latin American countries, has shaped recent Peruvian demography. For example, in 1961, only 50% of the 10 million Peruvians lived in urban centers, hereafter urban individuals, which has now risen to 79.3% ^24,25^. As in other parts of the world, the processes of admixture and urbanization overlap, involving complex socio-demographic phenomena with colonialist sequelae, which impact the genetic structure of human populations. Within this context, our understanding of how the interaction of continental ancestries (i.e., admixture dynamics) and the impact of internal migration shaped present-day Peruvian diversity is limited.

In this study, we address two questions regarding the genetic ancestries in Peruvians: (i) How diverse is the genetic composition, and how are their ancestries distributed across Peruvian urban areas? and (ii) how were the admixture processes between continental and regional groups in Peru during and after colonial times? To address these questions, we sampled and genotyped 13 urban Peruvian populations as part of the Peruvian Genome Project, including 346 newly genotyped samples and 504 individuals (including 418 Native American and 86 urban individuals) from recent studies ^8,9^ (Figure 1 and Table S1). A notable strength of our study is the sampling of predominantly Indigenous American populations. with new insights into ancestry composition, ancestry dynamics, and migration signature during the past ten generations. This work represents an effort to build resources to address genomic health equity.

**Figure 1.**
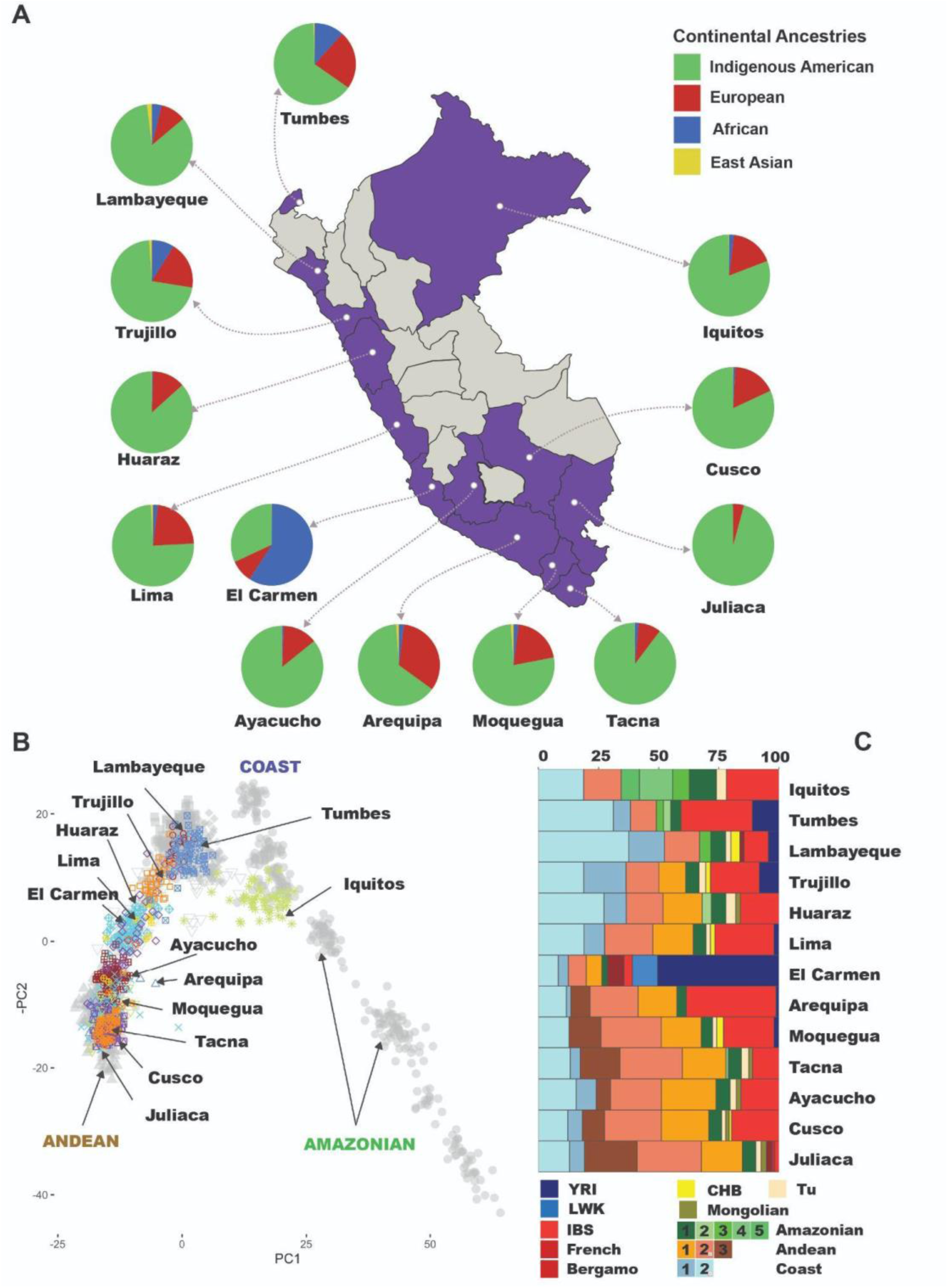
Ancestry composition in 13 urban Peruvian areas. A) Genome-wide ancestry proportions inferred by ADMIXTURE K=4 in urban centers of different dimensions, from Moquegua city (around 96,000 inhabitants) to Trujillo (around 970,000 inhabitants) and Lima, the Peruvian capital (around 10 million inhabitants). B) Ancestry-specific PCA for Indigenous American ancestries. Indigenous American references include populations from the Peruvian Pacific Coast, Andean, and Amazonian regions shown as grey circles. C) SOURCEFIND results for 13 admixed populations show a fine-scale inference of their ancestry proportions (See Table S2). For Indigenous American contributions, each number represents a population: Amazonian (1: Ashaninkas; 2: Awajun; 3: Candoshi; 4: Lamas; 5: Shipibo), Andean (1: Chopccas; 2: Qeros, 3: Uros), and Coast (1: Moche; 2: Tallanes). Indigenous American groups were described in Borda et al. (2020).

## Results

### About Sampling

Full details about sampling are described elsewhere ^26^. As part of the Peruvian Genome Project (PGP), 13 urban populations spanning diverse biogeographical regions were included (Figure 1). In this paper, we call urban centers a broad demographic spectrum, from small cities such as Moquegua with ∼96,000 inhabitants to Lima, a metropolis of around 10 million inhabitants. We collected blood from 643 individuals, none of whom were self-declared as indigenous. A subset of 346 samples was selected for genotyping using the IlluminaOmni2.5 array (<u>Table S1</u>). After quality control, our working dataset includes 2,034,418 SNVs. All procedures were evaluated and approved by the Peruvian Research and Ethics Committee of the Instituto Nacional de Salud (authorization no. OI-003-11 and no. OI-087-13).

### Fine-Scale Structure of Indigenous and Global Ancestries in Peruvian Genomes

We merged our dataset with the 1000 Genomes Project high coverage release ^27^, the Human Genome Diversity Project high coverage ^28^ datasets, and recently published PGP data^8,9^ (Table S2). This later dataset includes Indigenous American populations from the Peruvian Pacific Coast, Andean, and Amazonian regions. Genetic clustering analysis^29^ shows urban individuals as predominantly three-way admixed with a higher proportion of Native American ancestries (>60% [except for Afro-Peruvians], Figure 1A, Fig S1, Table S3) than European and African ancestries. In general, European and African admixture in urban populations is larger than in Indigenous populations ^8,9^ (Fig S1). Thirteen individuals showed East Asian ancestries (2-18%, Fig S1), likely as a result of the immigration of Chinese and Japanese individuals to Peru during the last 150 years, in part to replace the labor force in Coastal farms after the abolition of slavery in Peru in 1854 ^20,21^.

Our geographically diverse sampling allows us to identify the heterogeneity of ancestries in urban individuals. **ADMIXTURE (Fig S1) and principal component analyses** (PCA, Fig S2) revealed a latitudinal differentiation of Indigenous American ancestry proportions. Indigenous ancestry clusters related to the Pacific Coast and Amazonian Natives are predominant in northern urban populations, and Andean components are predominant in southern urban populations.

By coupling local ancestry analyses with PCA ^30,31^ (Ancestry-specific PCA: **asPCA**) restricted to Indigenous American segments, we showed that most urban groups clustered with Pacific Coast and Andean natives (Figure 1B). Interestingly, Southern urban groups (i.e., Ayacucho, Arequipa, Moquegua, Tacna, Cusco, and Juliaca) clustered with the Andean Native groups and showed more homogeneity compared with the Northern urban populations and Lima. We tested this homogeneity by performing **F_ST_ and Identity-by-Descent (IBD)** sharing analyses. We showed lower differentiation (lower F_ST_ values, Fig S3) and high-IBD sharing between Southern urban populations compared to Northern (Fig S4).

Furthermore, by exploring the haplotype sharing information using the **ChromoPainter-SOURCEFIND pipeline**^2,32^, we observed a different composition of Andean ancestries in urban Peruvian populations (Figure 1C, Table S4). Northern populations have two Andean components compared to the South, which includes a third Andean component associated with present-day Aymara speakers’ ancestry (Uros population). Consistent with ADMIXTURE results, Pacific Coast and Amazonian ancestries are more common in urban populations from Northern Peru. Also, SOURCEFIND suggests that the genetic composition of urban individuals can be explained as a mosaic of different Indigenous American ancestries.

European ancestry is the second most common continental ancestry in urban individuals (Figure 1A). During colonial times, the primary source of European ancestry was from Spain. However, after independence, the Peruvian government implemented policies that favored the migration of European communities to Peru ^33^. Our asPCA for European ancestries showed admixed Peruvians with higher affinity to a cluster that includes individuals from Spain (IBS) and Italy (TSI) (Fig S5). SOURCEFIND results confirm the predominance of South European ancestry in admixed Peruvians (Figure 1C, Table S4).

There is no precise number of how many enslaved Africans were introduced in Peru throughout the tragic transatlantic slave trade. Still, they concentrated in the coastal region^34^. Our ADMIXTURE and PCA results (Figure 1A, Fig S1) show most individuals with genome-wide African ancestries in the coast region, mainly in the North (Trujillo and Tumbes) and South Coast (El Carmen). During the initial decades of the colony, the majority of enslaved Africans originated from West Africa, specifically between the Senegal and Niger rivers^34^. However, by the late 1580s, the Congo-Angola region (Central-Southern Africa) emerged as a primary source^34^. We explored the diversity of African ancestries in admixed individuals by performing asPCA and SOURCEFIND. In the asPCA, we observed admixed Peruvians with higher similarity to present-day West-Central African populations but also some genetic affinity to East African (LWK) populations, a population group with more genetic affinity to other Central-Southern African individuals. This pattern would suggest some admixture between these components. This is supported by SOURCEFIND results, in which the African component is higher in Northern populations and El Carmen (> 10%, Table S4) and is shown as admixed from West-Central and Eastern African related ancestries (Figure 1C, Table S4).

It was after Peruvian independence that slavery abolition was achieved^35^. However, driven by economic interests, an alternative source of inexpensive labor was sought. This led to the influx of East Asian migrants into Peru, with approximately 100,000 Chinese individuals arriving in the 19th century^20^. The ADMIXTURE results revealed minimal individuals with East Asian-related ancestries (Figure 1A). Our asPCA analysis of East Asian ancestries indicated that most individuals clustered with the Southern Chinese population, aligning with historical records. Notably, two individuals from Southern Peru (Moquegua) exhibited a higher affinity with Japanese individuals. At the population level, SOURCEFIND detected more East Asian ancestries (∼6%) in Lambayeque, a city on the Peruvian Coast (Figure 1C and Table S2).

### The signature of admixture shaped Present-day admixed Peruvians

Our second question investigated how continental and regional groups interacted in Peru during and after colonial times. To address this, we applied two approaches, GLOBETROTTER^36^ and *HierarchyMix*^37^. GLOBETROTTER examines LD decay observed on co-ancestry curves, providing the timing and mode of admixture events and ancestry composition for the hypothetical sources. In contrast, *HierarchyMix* analyzes the distribution of local ancestry segments. While GLOBETROTTER provides a subcontinental perspective on admixture events, *HierarchyMix* focuses on the interaction of continental ancestries.

We inferred two major admixture events (See Methods, Figure 2, Fig S6-S8). GLOBETROTTER detected a single admixture event, but the admixed profile of its sources suggests an earlier admixture event (Figure 2). Complementarily, *HierarchyMix* inferred admixture events per each pair of continental ancestries. Except for El Carmen, recent events for all Peruvian populations were inferred around 8-12 generations ago, or within the past 215-322.8 years (Figure 2), assuming a generation time of 26.9 years^38^. Most populations showed these events as the admixture between a predominantly indigenous American source and a non-indigenous source (Figure 2, Fig S6-8). Both methods inferred the non-indigenous source as admixed of European and African ancestries, except for Trujillo and El Carmen. *HierarchyMix* dated this earlier events of African-European admixture around 12-25 generations (Fig S6-8). Moreover, for Trujillo and El Carmen, the earlier admixture event involved Europeans and Indigenous Americans, followed by a recent African contribution (Figure 2, Fig S6-8).

**Figure 2.**
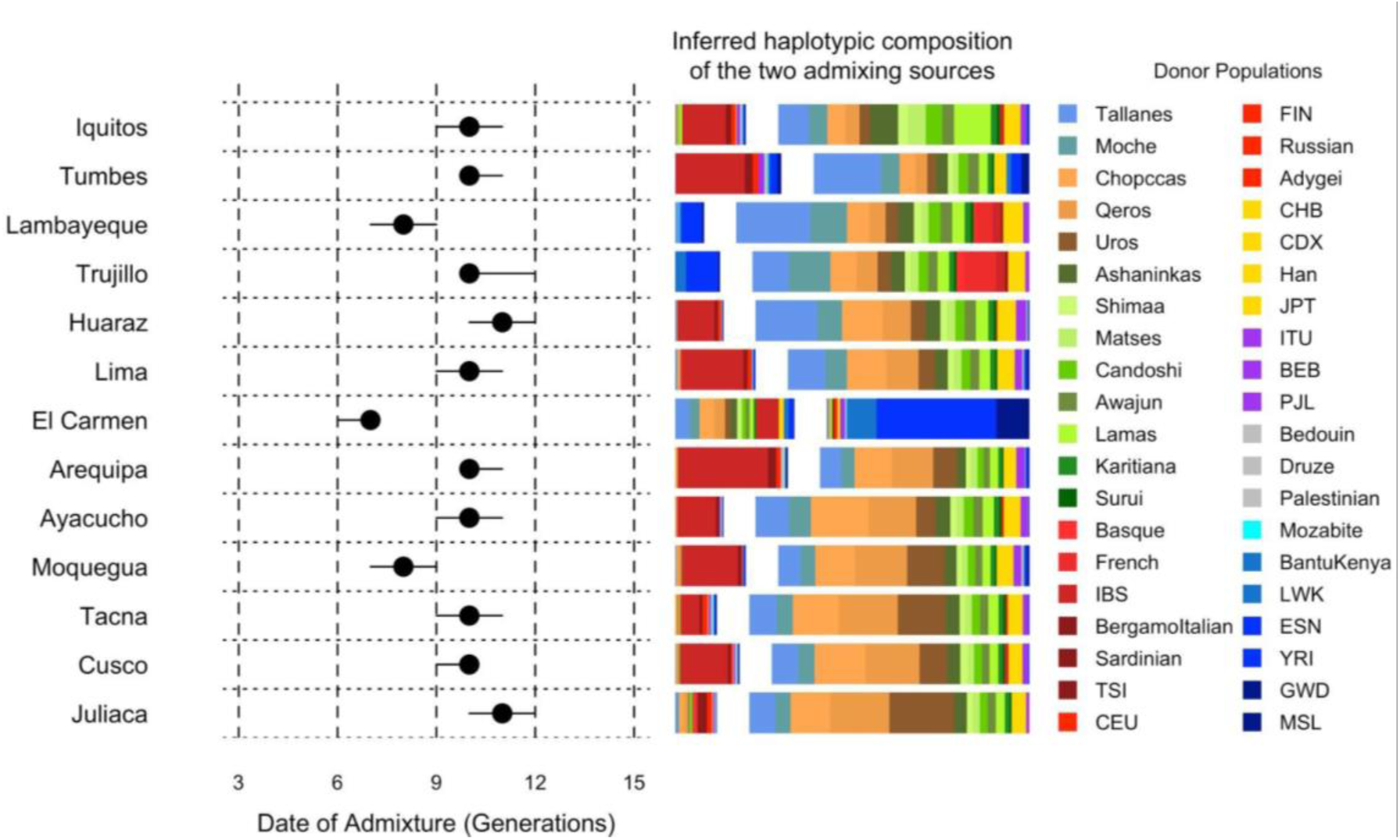
Admixture dynamics and recent gene flow in 13 urban Peruvian populations. GLOBETROTTER results for admixture timing and mode in 13 urban Peruvian individuals. The left panel shows the inference of admixture timing with the point estimate of the admixture date shown as a black point, with a 95% confidence interval shown with lines. Bar plots represent the admixture sources inferred by GLOBETROTTER, colored by donor populations as in the legend. The population order from the top to the bottom follows a north-to-south pattern.

The subcontinental composition of the Indigenous source inferred by GLOBETROTTER aligns with SOURCEFIND results (Figure 1C), where Andean ancestries dominate in southern groups. This pattern reinforces that the present-day structure in the admixed population mirrors the Native American structure of the last 2000 years^14^. Moreover, the admixed genetic profile of the sources supports older admixture events and more complex dynamics between indigenous groups during the genetic history of Peruvians, as described in Harris et al^8^.

*HierarchyMix* also inferred admixture events involving East Asian and Indigenous American ancestries. These events were dated several generations before the historical arrival of East Asian migrants in the 19th century, and considering the small contribution of less than 1% in most cases, this signal could be a spurious result from local ancestry inferences. However, our SOURCEFIND results support a significant East Asian contribution to the Peruvian population, with Lambayeque with the highest proportion (∼6%). The predominantly Indigenous source for the recent admixture event inferred by GLOBETROTTER showed a small proportion of East Asian ancestries in northern populations (Figure 2). However, it is unclear if this is confounding and represents the genetic composition of unsampled Indigenous American ancestries.

### Reconstructing Post-colonial demographic events

The GLOBETROTTER and *HierarchyMix* results broadly overview major admixture events involving continental ancestries around nine to 12 generations. Considering this, we assumed that DNA shared during the last eight generations could be associated with gene flow among urban groups. Our population structure analyses detected a higher genetic similarity among southern populations (Figure 1B). Is this higher similarity the result of ancient gene flow during pre-colonial times, recent gene flow during post-colonial times, or both?

To address this question, we investigated patterns of gene flow through IBD segments shorter and longer than 18.75 cM. These two intervals explain the DNA sharing due to the recent common ancestor of two individuals before and after approximately eight generations ago ^39^, respectively (See Methods). The short segment interval shows increasing IBD sharing from North to South (Figure 3A). This mirrors the full IBD sharing patterns (Fig. S4), underscoring the influence of ancient gene flow (pre-colonial times) and admixture (colonial times) on increasing the genetic similarity in Southern Peru. Importantly, this pattern has seemingly continued for the last eight generations (Figure 3B), indicating that recent gene flow follows ancient trends. Furthermore, high IBD sharing between Trujillo and Afrodescendants from El Carmen potentially reflects shared African ancestry. In summary, ancient and recent gene flow have driven the reduced differentiation, or increased connectivity, observed in populations from Southern Peru compared to northern populations.

**Figure 3.**
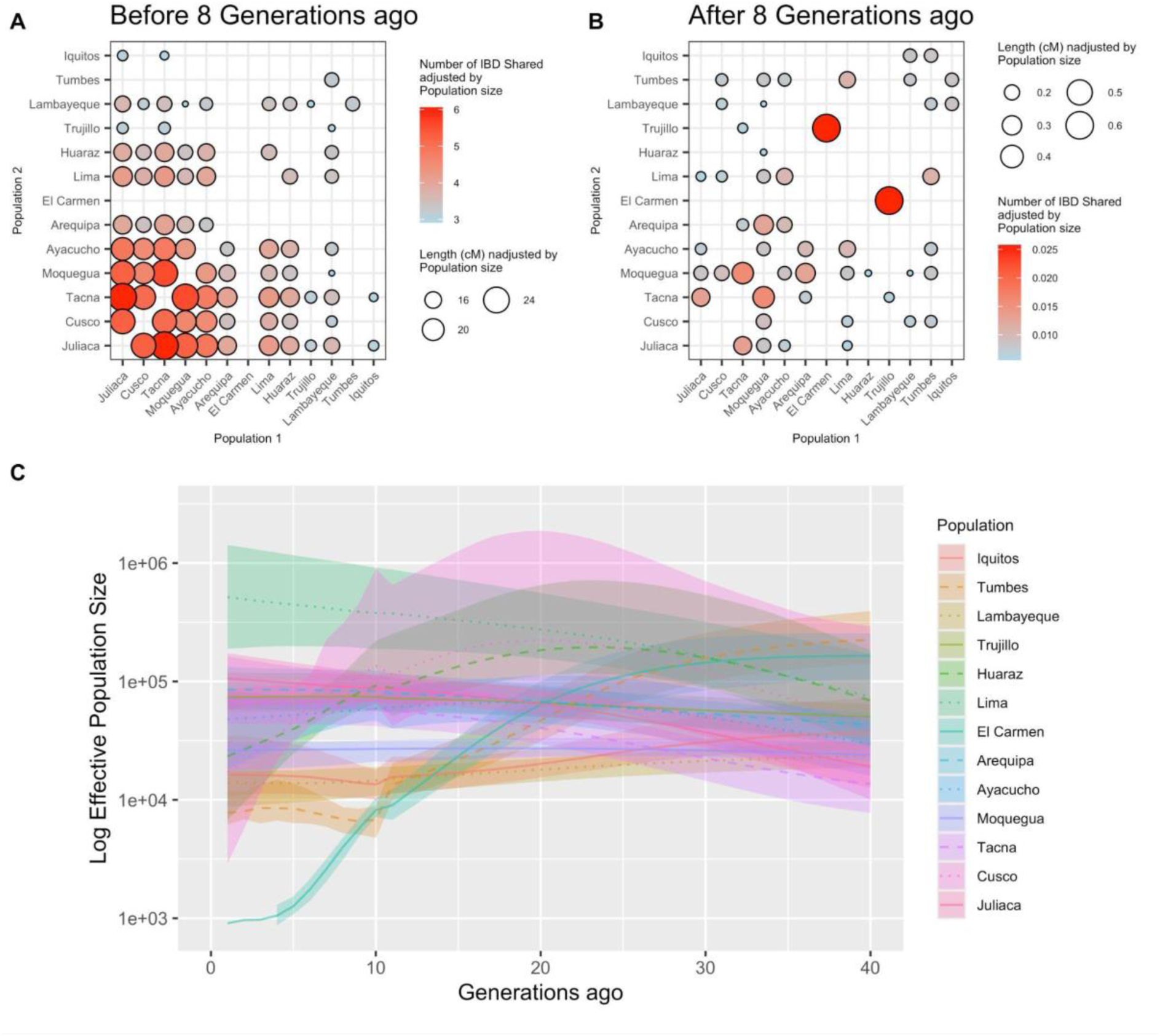
Demographic patterns between 13 urban Peruvian Populations. We organize IBD segments in two intervals corresponding to (A) pre- and colonial gene flow between 13 urban Peruvian populations and (B) Postcolonial and post-admixture events. The first interval includes IBD segments smaller than 18.75 cM. This IBD interval represents the DNA sharing of the last eight generations. For visualization purposes, internal sharing was ignored. The population order from the top to the bottom follows a north-to-south pattern. C) Effective population size inferred through HapNe for 13 urban populations. Lima (green shade with dotted line) shows continuous population growth.

Furthermore, we leveraged the IBD segment distribution to estimate the trajectory of the effective population size (*Ne*) in recent generations using HapNe^40^ (See Methods). Our analyses revealed a notable population growth in several southern urban groups, including Tacna, Arequipa, and Juliaca, compared to northern groups (Figure 3C, Fig S9-S10). As the capital, Lima notably shows an even steeper increase in *Ne*, consistent with recent migration trends toward the city and with the fact that it includes a third part of the country’s total population. In contrast, the El Carmen population, predominantly self-identified as Afro-Peruvian, displays a declining *Ne* trajectory over the past 25 generations, underscoring a distinct demographic pattern, likely characterized by isolation.

The colonial admixture was characterized by the encounter of continental ancestries, leaving a signature of sex-bias admixture reported in several Latin American populations ^41,42^. Considering our fine-scale sampling, we evaluated sex bias admixture and its regional differences. A comparative analysis of ancestry proportions revealed that, in nine of thirteen populations, European ancestry was significantly higher in autosomes compared to the X chromosome (Wilcoxon test, p < 0.05; Fig S11, Table S3). The complementary pattern was observed for Indigenous American ancestry (Fig S11 and Table S3). The African ancestries showed a similar pattern to that of Europeans in five populations, but with small proportions (Fig S11 and Table S3). These results suggest that, with some exceptions, European admixture in Peruvians was preferentially mediated by males.

## Discussion

Admixed populations in western South America exhibit high levels of Indigenous American ancestry ^8,43^, contrasting with the pattern seen in other South American groups ^2,44–47^. This manuscript, focused on admixed populations from Peru, builds upon the first phase of the Peruvian Genome Project ^8,9,48^ and incorporates new genome-wide data. Through our manuscript, we replicate previous results and present three key new findings about Peruvian genetic composition: (i) The population structure of urban individuals reflects a North-South gradient akin to that observed among Indigenous Peruvians, owing to their elevated proportion of Indigenous American ancestries; (ii) Most continental admixture events before eight generations ago were followed by internal migration patterns that align with the North-South continuum; and (iii) the trajectory of the effective population growth in Lima reflects the recent process of migration from rural to urban pattern.

Multiple disciplines, including archaeology, linguistics, and genetics, revealed that geography shaped the population structure of Indigenous groups from western South America into two ancient spheres of cultural interaction ^9,14,15^, a Northern and a Southern. This is consistent with an autosomal STR study of 20 of the 24 Peruvian administrative regions that identified Peruvian population structure following a latitudinal rather than longitudinal structure ^13^. Our primary contribution lies in elucidating the distribution of subcontinental ancestries along the latitudinal axis. For example, our Indigenous American panel included Andean populations from two linguistic families, Quechua and Aymara speakers. We demonstrate that ancestry similar to present-day Aymara speakers is more common in the South, where most Aymaran Indigenous groups live. At the same time, the Quechua-related similarity was inferred in all Peruvian samples in this study, consistent with the wide geographical distribution of Quechua speakers in the Andean region that resulted from interactions spanning the past 2000 years, including Pre-Inca, Inca, and colonial migrations ^14,49,50^.

Our second contribution explains the dates and mechanisms through which ancestral source populations interacted, shaping the present-day genetic composition of an urban area population. Here, we identified a complex admixture pattern that showed at least two admixture events. The most recent admixture events were during the last eight to ten generations, but the ancestry composition of the sources of these events suggests previous admixture events. Notably, both methods revealed African-European or Indigenous American-European interactions, but no African-Indigenous American. During the early colonial period, Spaniard invaders imposed social construction that divided enslaved Africans and Indigenous American people ^19^, leading to conflicting interactions.

Regarding the contribution labels, the inferred European-related sources in the genetic pool of admixed Peruvians align with historical records, as expected, indicating significant contributions from Southern Europe ^49^. For African ancestries, our results suggest some admixture between components associated with West, West-Central, and East African regions. Our GLOBETROTTER analyses indicate that these ancestries were admixed before the last ten generations. One inferred ancestral source shows the earlier event for Afrodescendants from El Carmen (Figure 2), showing these three components. By the time of independence in 1821, Spaniard invaders had imported over 100,000 enslaved African people to Peru to meet agricultural demands, mainly on the North Coast during the Transatlantic Trafficking of Enslaved Africans ^50,51^. They were brought from ports in the Caribbean region. In this region, the European invaders mixed African individuals from different ethnicities to sow confusion and prevent an uprising.

For East Asian ancestries, their distribution and diversity align with historical records. We observed these ancestries among individuals in coastal and northern Peru (Figure 1C). In these regions, economic activities were undertaken by East Asian migrants following the abolition of slavery, albeit under conditions as dire as slavery itself ^20^, since it played a pivotal role in the economic advancement of colonies. However, given the small proportion of ancestry, we have insufficient power to accurately detect the correct admixture date for this component.

Our third contribution highlights major demographic events inferred through IBD analyses. The asPCA and Fst results revealed higher genetic similarities among southern urban areas. We examined the distribution of IBD sharing intervals corresponding to the past eight generations to investigate whether this pattern was driven solely by ancient migrations or by a combination of ancient and postcolonial movements. Our findings indicate that genetic similarities among urban areas in southern Peru are not only due to ancient patterns observed in Indigenous Americans from the region but also reflect continuity in migration patterns over the last ten generations. Moreover, IBD inference helps to explain the urbanization process in Peruvian groups. Lima, the capital of Peru, has long attracted migrants from other regions due to its disproportionate economic development compared to other Peruvian cities. Over the years, thousands have relocated there, making Lima’s population ten times larger than major cities like Trujillo, Arequipa, and Cusco ^52^. This trend reflects broader urban migration driven by socioeconomic disparities and political violence, even in recent decades ^53–55^. By analyzing the effective population size evolution inferred from IBD segments, we recapitulated this growing pattern, which has the consequence of increasing its genetic diversity.

Three limitations can be observed in our study: (a) diversity gaps, poor sampling of the central and Southern Amazonian regions,(b) small sample size to perform demographic analyses, and (c) the power of haplotype-based methods to discriminate ancestries. Our first limitation does not allow us to explore the three clusters identified by Innaconne et al. ^13^. We expect more genomic initiatives to explore the genetic diversity of Indigenous and Admixed groups in central Peru, an important area where the Andean and Amazonian biogeographic regions interact, providing ecosystems with particular selective pressure that shapes their phenotypes. However, our sampling provided interesting clues regarding the diversity and distribution of ancestry components. Our second limitation is our lack of statistical power to correctly identify demographic events related to admixture dynamics and effective population size. For example, our IBD sharing analysis relies on observed IBD distribution, which could be biased due to phasing error, inaccurate IBD detection, or sampling not reflecting population-level patterns. These issues can affect the inference of effective population size; however, our interpretation is based more on the complete picture of the distribution than on particular details. Our third limitation concerns the power of local ancestry inference to discriminate between continental ancestries. This is the case for the inference of segments with East Asian origin, which could be misclassifications of Indigenous American origin and vice versa, generating conflicting results for the inference of admixture between these two ancestral groups.

Overall, this work represents another important step in studying genetic history in admixed groups with predominantly Indigenous American ancestry. Several efforts have been performed in Latin America^9,10,44,45,56^, which, together with our study, represent the effort of local institutions to understand the diversity of their people, people who were underrepresented, but an entire world of genetic and epidemiological histories that we need to understand.

## Acknowledgments

We are deeply grateful to all participants who generously donated blood samples for the Peruvian Genome Project. Their contribution was vital to the success of this research. We thank all those who facilitated the recruitment of participants, including the Direcciones Regionales de Salud from Loreto, Puno, Cusco, La Libertad, Huancavelica, Ica, Piura, Ancash, Arequipa, Ayacucho, Tacna, Ucayali, San Martin, Amazonas; Universidad Andina Nestor Caceres Velasquez Facultad de Ciencias de la Salud, Universidad Nacional Jorge Basadre Grohmann, Universidad Nacional Mayor de San Marcos, Universidad Nacional San Agustín, Universidad Nacional de San Cristóbal de Huamanga, Universidad Nacional Santiago Antúnez de Mayolo, and Universidad Nacional de Trujillo. This work was supported by the University of Maryland Institute for Health Computing (V.B. and B.A.M.), the Institute for Genome Sciences and Program in Personalized Genomic Medicine at the University of Maryland School of Medicine (V.B., D.V.O., and T.D.O.), and the Instituto Nacional de Salud, Lima, Perú (O.C., C.S., C.P., and H.G.). ET-S and CS-C are supported by Conselho Nacional de Desenvolvimento Científico e Tecnológico (CNPq), Fundação de Apoio à Pesquisa do Estado de Minas Gerais (FAPEMIG), Coordenação de Aperfeiçoamento de Pessoal de Nível Superior (CAPES), and the Brazilian Ministry of Health (Department of Science and Technology-DECIT). M.M was supported by the CGEn HostSeq/CIHR fellowship (CGE 185054) and the SickKids Restracomp Fellowship.

## Contributions

The project was conceived by V.B, O.C., C.S., C.P., E.T.S, H.G., and T.D.O. Sampling and Genotyping data was performed by O.C., C.S., C. P., and H.G. Quality control was performed by V.B. Data analyses was performed by V. B., D.V.O.,M. M., C.S.C., I.A., B. A. M., and M.H.G. All authors contributed with data interpretation. V.B., E.T.S, and T.D.O. wrote the manuscript. All authors read the manuscript and contributed with suggestions.

## Competing interests

The authors declare no competing interests.

## Methods

### Sampling, Genotyping, and Quality control

Thirteen populations were sampled in 2010. All details related to sampling were described elsewhere^26^. Samples were genotyped using the Illumina Omni2.5M array in several batches at the Instituto Nacional de Salud (Peruvian National Institute of Health) facilities. We exported raw data to PLINK format using GenomeStudio. We used PLINK1.9^57^ to perform all quality control steps for autosomes and chromosome X. The quality control process at the single-nucleotide variant (SNV) level included the following steps:

- Exclusion of variants with no chromosome information
- Exclusion of AT/CG genotypes
- Exclusion of variants with 100 Heterozygous
- SNVs out of the Hardy-Weinberg equilibrium (HWE) (threshold 10e-6)

Some batches were genotyped using GRch37. For these batches, liftover to GRch38 was performed using the UCSC liftover tool (https://genome.ucsc.edu/cgi-bin/hgLiftOver) before merging with other GRch38 batches. After merging, we filtered for SNPs and individuals accumulating more than 5% of missing data using PLINK (--geno 0.05 and mind 0.05).

We controlled for relatedness using the KING algorithm^58^ to identify duplicates and individuals with first- and second-degree relationships. We ran the KING algorithm in the SNPRelate package ^59^ using the `snpgdsIBDKING` function. To decide which individuals should be removed after relatedness analyses, we used NATORA^60^. This admixed genotyped data was merged with Peruvian Indigenous American data previously reported^8,9^ to build the PGP dataset that included 749 individuals (Admixed and Indigenous American Peruvians) and 1,933,621 SNVs.

### Phasing

We set the reference allele to the GRCh38 reference allele using the `GRCh38_full_analysis_set_plus_decoy_hla.fa` file and PLINK. Following this, we phased PGP data using shapeit4^61^ with a reference panel that includes the full 1KGP High coverage ^27^ (n=3,199 individuals) and HGDP data ^28^(n=827 individuals). To improve phasing quality, we set the mcmc iterations parameters to `10b,1p,1b,1p,1b,1p,1b,1p,10m.` After phasing, we merged Phased PGP with 1KGP and HGDP. Also, we removed multiallelic variants and AT/CG genotypes. Finally, our combined dataset included 4,775 individuals, 1,461,568 autosomal SNVs, and 25,418 chromosome X SNVs.

### Population structure analyses

We used allele frequency-based methods to explore population structure in our combined dataset. First, we generated a list of unlinked variants using the SNPRelate `snpgdsLDpruning` function with a maximum base pair of 1Mb for the sliding window, a correlation method to calculate LD values, and a threshold of 0.1 With KING relatedness inference and the unlinked SNP list, we ran PCAir^62^ with a minor allele frequency (MAF) threshold of 5%, kinship threshold of 0.044194, and divergence threshold of -0.044194.

To explore genome-wide ancestry proportions, we ran ADMIXTURE^29^ from K=4 to K=8 in a subset of individuals that includes Full PGP data (admixed and Native Americans), a subset of 1KGP high coverage populations with unrelated individuals (TSI, IBS, CEU, PEL, CLM, MXL, PUR, LWK, MSL, YRI, GWD, JPT, CHB, and CHS), and Native Americans from HGDP (Colombian, Pima, and Maya populations). This subset included unlinked SNVs, and an MAF filter of 5%. We performed ten runs per K value and plotted the run with the highest log-likelihood. To explore differentiation among urban populations, we ran Fst analysis on PLINK using the Weir and Cockerham estimator ^63^.

### Identity-by-Descent and Local Ancestry Inferences

We removed SNVs with MAF < 0.001 of the full phased data and inferred identity-by-descent (IBD) segments using hap-ibd ^64^ with default parameters. After IBD inference, we ran the merge-ibd-segments software to remove gaps between IBD segments. We kept IBD segments greater than 3cM for subsequent analyses and removed ROH segments.

We performed local ancestry to infer the continental ancestry at the chromosomal level. For this purpose, we ran GNOMIX^65^ with four continental references based on ADMIXTURE results (Population Structure analyses section). We selected European (n=404 individuals; Populations: CEU, GBR, IBS, and TSI), African (n=405 individuals; Populations: YRI, ESN, GWD, and LWK), and East Asian (n=411 individuals; Populations: CHB, CHS, JPT, and KHV) individuals from 1KGP populations. Native American reference included 187 individuals from PGP with more than 99% NAT ancestry based on ADMIXTURE K4 results. We ran GNOMIX using the option “best" for model inference parameters. This option uses random string kernel base + xgboost smoother and is recommended for array data. We also include the rephasing option to correct the phase inference based on local ancestry information.

We ran ancestry-specific PCA to explore the similarity between chromosomal ancestry segments of PGP individuals to specific continental ancestral references using the Browning et al. pipeline ^31^ (http://faculty.washington.edu/sguy/local_ancestry_pipeline/). Before running the asPCA, we convert msp GNOMIX output to viterbi output since the pipeline was designed for rfmix ver1 outputs. The msp output includes ancestry information for loci in each pair of haplotypes. Considering a 4-way admixture, we created an ancestry-specific msp file setting as missing the other three ancestries. To perform ASPCA, we kept individuals with a minimum of 30% for European or African ancestry. For East Asian ancestry, we kept individuals with a minimum of 5%.

### Admixture dynamics analyses

We explored admixture dynamics in Peruvians by analyzing different evolutionary aspects: 1) Admixture timing and mode, 2) Recent migration explored by IBD sharing, 3) Demographic changes identified by the evolution of effective population size (*Ne*), and 4) Sex-bias admixture analyses.

First, to analyze the admixture timing and mode, we ran GLOBETROTTER and HierarchyMix to explore admixture timing and mode.

> To run GLOBETROTTER, we first ran ChromoPainter-based methods^2,32,36^ to identify the timing and mode for admixture events. This analysis requires several steps: i) inference of scaling parameters [recombination and mutation rates], ii) inference of haplotype sharing (copyvectors and painting samples), iii) inference of ancestry proportions using haplotype information, and iv) inference of admixture date and mode.
>
> **i) Inference of scaling parameters**: We ran ChromoPainter version 2 (https://github.com/sahwa/ChromoPainterV2) to determine scaling parameters for haplotype sharing inference. For this purpose, we performed a first ChromoPainter run in a subset of chromosomes (1,8,15,22) and a subset of individuals to calculate the scaling parameters.
> **ii) Inference of haplotype sharing**: After calculating the scaling parameters, we set these values in two ChromoPainter runs including all individuals and chromosomes to generate two types of files: copyvectors and painting samples. Our second Chromopainter was to infer copyvectors, we set the parameter -f to select the donors and recipients. We selected all populations as recipients but just Indigenous Americans, European, African, and Asian populations as donors (Supplementary Table 2). Our third Chromopainter run was to infer painting samples, we set the -f mode setting admixed Peruvians as recipients and all other populations as donors.
> **iii) Ancestry proportions at the subcontinental level**: After ChromoPainter runs, we merged copyvectors into one coancestry matrix using the neaverage perl script (https://people.maths.bris.ac.uk/~madjl/finestructure/neaverage.pl.zip) and ran SOURCEFIND^2^ ver2 (https://github.com/hellenthal-group-UCL/sourcefindV2) to identify ancestry proportions corresponding to donor populations. SOURCEFIND is a Bayesian approach that explores haplotype sharing between donor individuals and recipients to determine ancestry proportions beyond the continental level.
> **iv) Admixture inference:** To determine the date and mode of admixture events, we ran GLOBETROTTER^36^ (https://github.com/hellenthal-group-UCL/GLOBETROTTER), a software that explores ancestry proportions and linkage disequilibrium decay. We ran GLOBETROTTER in each population and performed 100 bootstrap replications. Crucially, in assessing whether the inferred admixture event in a population signifies a “one-date” or “multiple-dates”, we checked the ancestry curves and the goodness-of-fit (R2) for a single date of admixture (maxR2fit.1date). We determined that if the maxR2fit.1date exceeded 99.9% (as communicated by Garret Hellenthal personally), we interpreted the result as indicative of a one-date admixture, adopting the date and proportions proposed by this mode. However, it’s important to note that this inference pertains to the most recent event and does not preclude the possibility of earlier events.
>
> For *HierarchyMix* Inference, we use GNOMIX outputs in bed format to run *HierarchyMix* with default parameters except the number of bootstrap replicates and threshold for ancestry segments. We ran 1000 bootstrap replicates and kept segments greater than 2cM.

Second, we analyzed IBD sharing among PGP populations to explore recent migration. We kept IBD segments greater than 18.75cM to explore sharing among populations during the last eight generations that resemble Colonial and post-colonial events. Considering populations X, Y, and Z, we assumed that any IBD segment shared among X and Y populations corresponds to people’s movement among them. However, we cannot rule out that sharing segments between populations could also explain migration from a third source (population Z) to both populations (X and Y) during the same period. To estimate the interpopulation sharing after keeping IBD segments greater than 18.75 cM, we sum the number and the length of IBD segments shared between individuals from different populations. To control for sample sizes, both, the total number and the total length were divided by the product of the sample size of both populations.

For plotting IBD and GLOBETROTTER results, we modified R codes available at:

- https://github.com/georgebusby/admixture_in_africa
- https://github.com/chiarabarbieri/SNPs_HumanOrigins_Recipes

Third, we inferred the distribution of the effective population size during the last 50 generations for PGP data using HapNe^40^ following the pipeline described here: https://github.com/PalamaraLab/HapNe.

Briefly, for each population, each phased chromosome was split into two parts, each divided by the centromere using PLINK. Then, each chromosomal segment was used to infer IBD segments using hap-ibd and then merged with the merge-ibd-segments.jar software. Finally, these outputs were used as inputs for HapNe with default parameters.

Fourth, to interrogate whether a population resulted from sex-bias admixture, we compared admixture proportions from autosomes and chromosome X. For autosomes, we used results from the ADMIXTURE analyses described above. To infer continental ancestry proportions for chromosome X, we ran ADMIXTURE using the --haploid flag. We compared autosomal and chromosome X proportions using the Wilcoxon test.

### Resources

GRCh38 fasta file: https://ftp-trace.ncbi.nih.gov/1000genomes/ftp/technical/reference/GRCh38_reference_genome/GRCh38_full_analysis_set_plus_decoy_hla.fa

shapeit4: https://github.com/odelaneau/shapeit4/tree/master

ASPCA: https://faculty.washington.edu/sguy/local_ancestry_pipeline/

hap-ibd: https://github.com/browning-lab/hap-ibd

merge-ibd-segments: https://faculty.washington.edu/browning/refined-ibd/merge-ibd-segments.17Jan20.102.jar

HapNe: https://github.com/PalamaraLab/HapNe

ChromoPainter: https://github.com/hellenthal-group-UCL/ChromoPainter

SOURCEFIND: https://github.com/hellenthal-group-UCL/sourcefindV2

GLOBETROTTER: https://github.com/hellenthal-group-UCL/GLOBETROTTER

HierarchyMix: https://github.com/Shuhua-Group/HierarchyMix

## Data and code availability

Data will be deposited at the European Genome-phenome Archive (EGA), which is hosted by the EBI and the CRG. The accession number will be available soon. Further information about EGA can be found on https://ega-archive.org "The European Genome-phenome Archive of human data consented for biomedical research" (https://doi.org/10.1093/nar/gkab1059).

## Supplementary Figures

**Figure S1.**
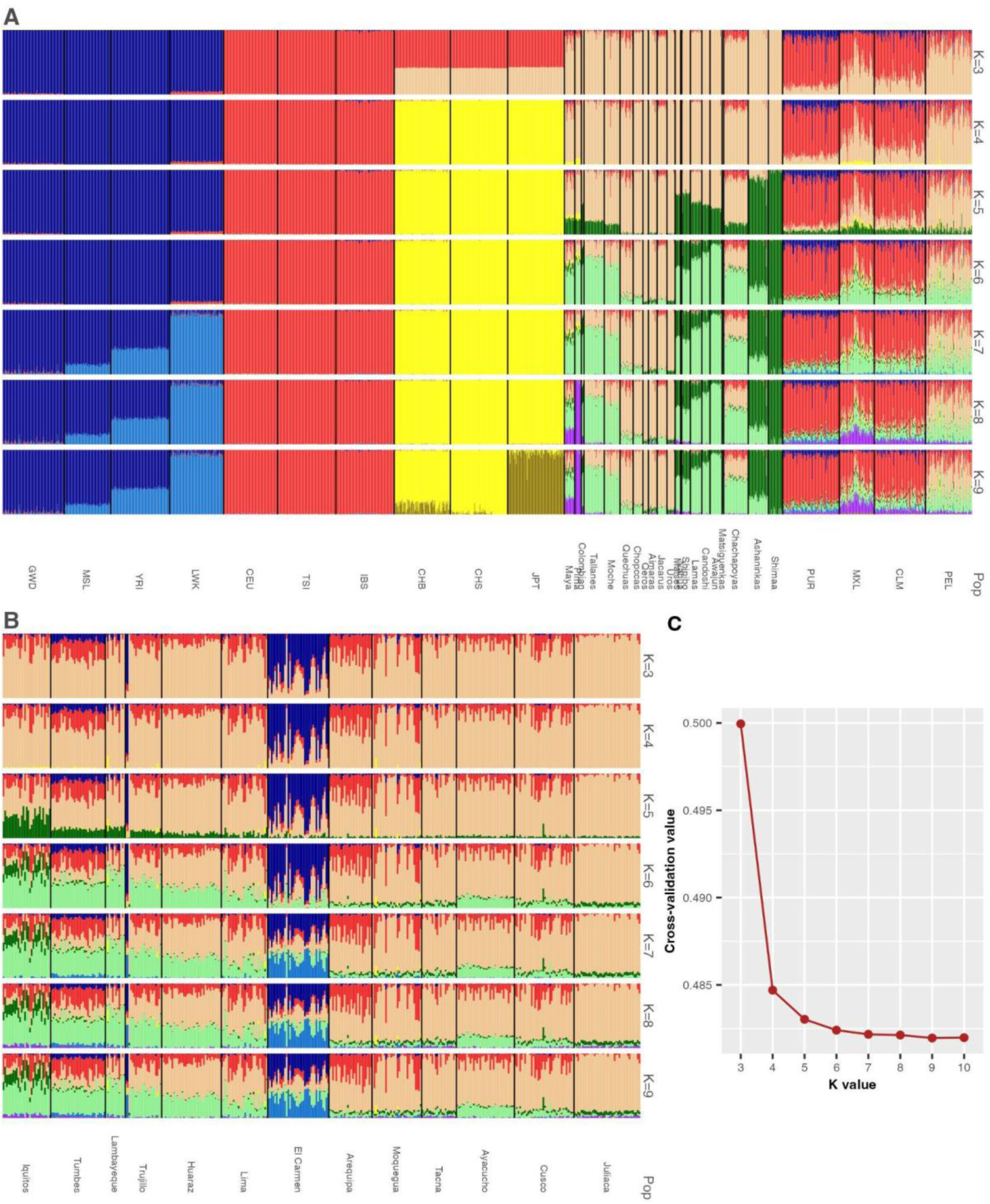
Genetic clustering analyses to infer ancestry clusters in urban individuals from the Peruvian Genome Project (PGP) and Reference populations. ADMIXTURE was run on unsupervised mode from K=3 to 9 for the dataset, including urban populations from Peru, Native Americans from PGP (See Borda et al. 2020^9^), and HGDP^28^ (Colombians, Maya, and Pima), and European, African, and East Asians from 1000 Genomes Project ^27^. All these populations were included in each run. A) ADMIXTURE plots of reference populations for K values from 3 to 9. B) ADMIXTURE plots of urban populations for K values from 3 to 9. C) Cross-validation values for runs from K=3 to K =9. Pop = Population.

**Figure S2.**
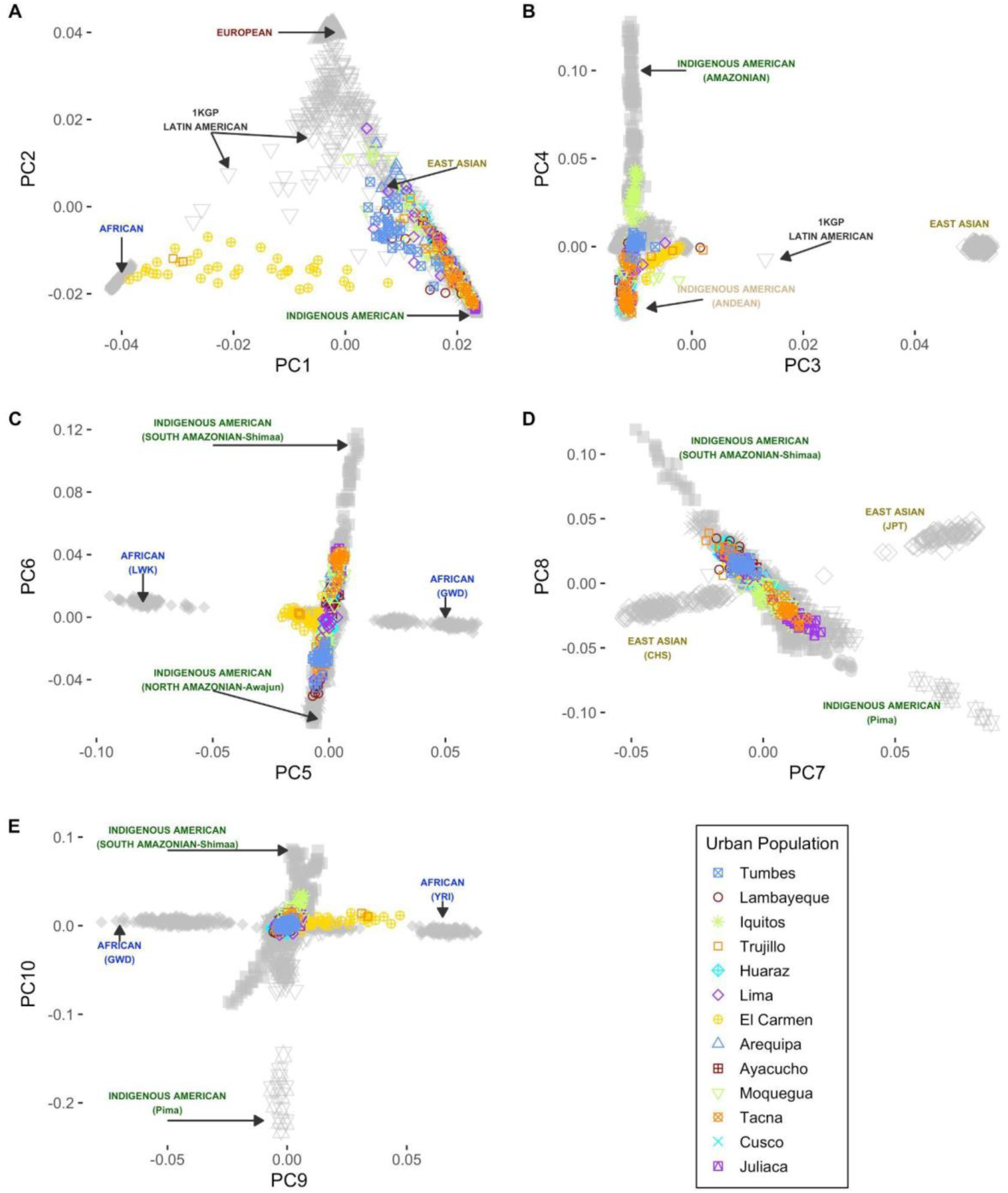
Principal Component analyses for Peruvian admixed individuals in a worldwide context. We run PCA using PCAir for a dataset that includes PGP data and 1000 Genomes Project + HGDP populations.

**Figure S3.**
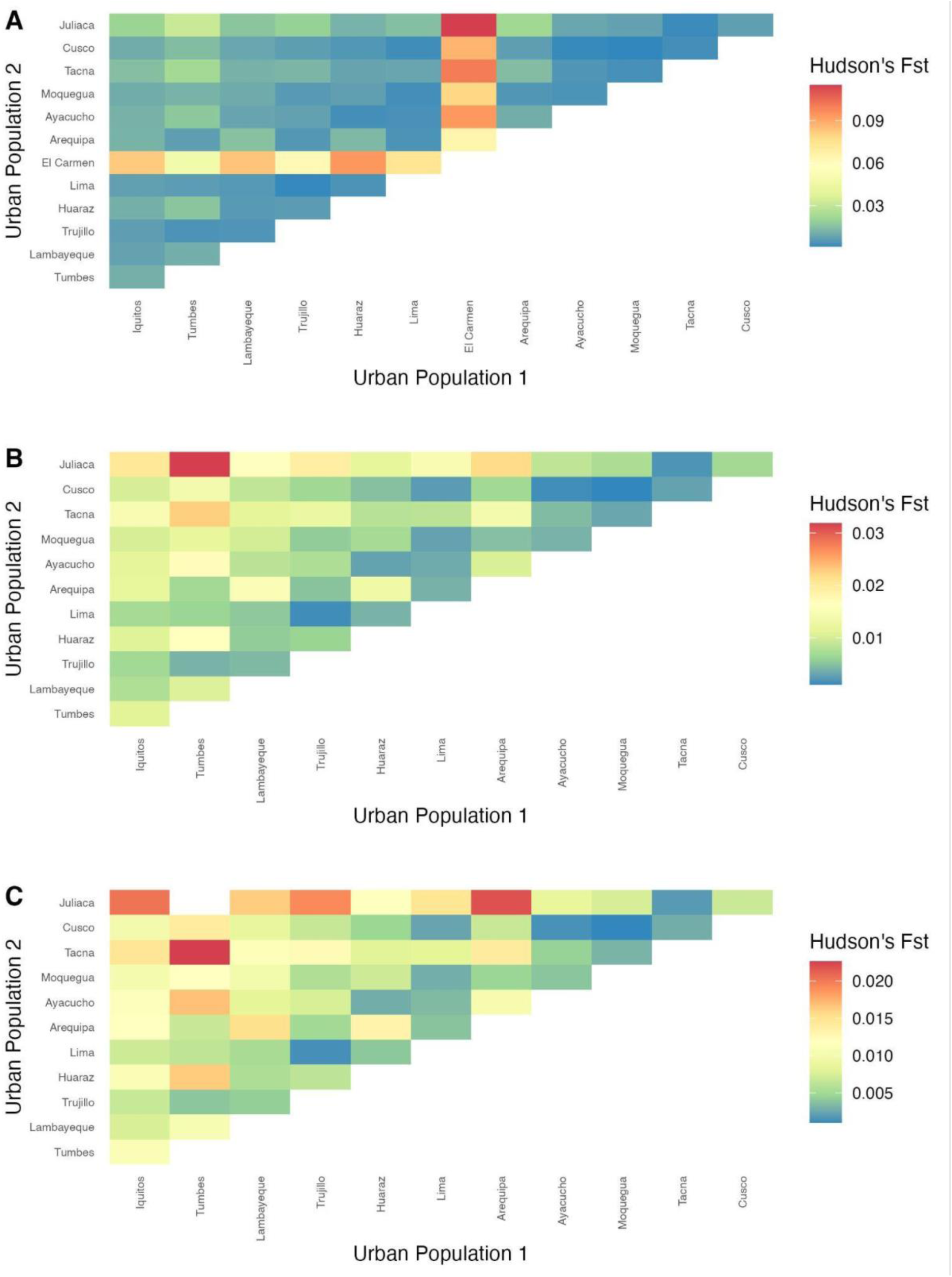
FST results. A) All populations, B) All populations except Ica (Afrodescendants), C) Removing Fst between Puno (the southernmost part of Peru) and Tumbes (the northernmost part of Peru).

**Figure S4.**
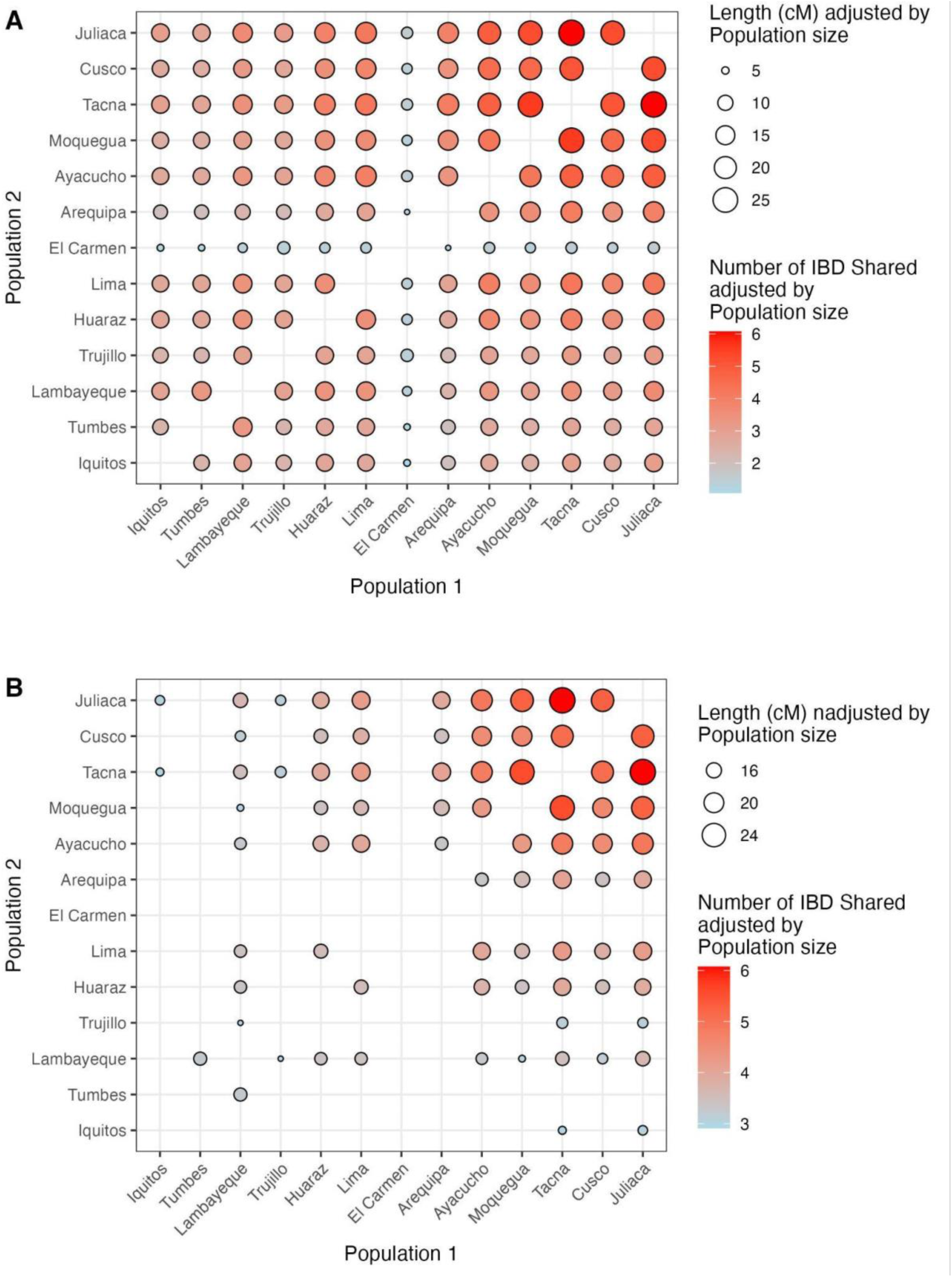
Identity-by-descent (IBD) sharing among Peruvian admixed populations. We included greater than 3cM and removed the intrapopulation sharing to explore gene flow and relatedness among populations. The total length and number of IBD segments were adjusted by population size. A) Total IBD sharing for all populations. b) Total IBD sharing for all populations with a sharing higher than the median.

**Figure S5.**
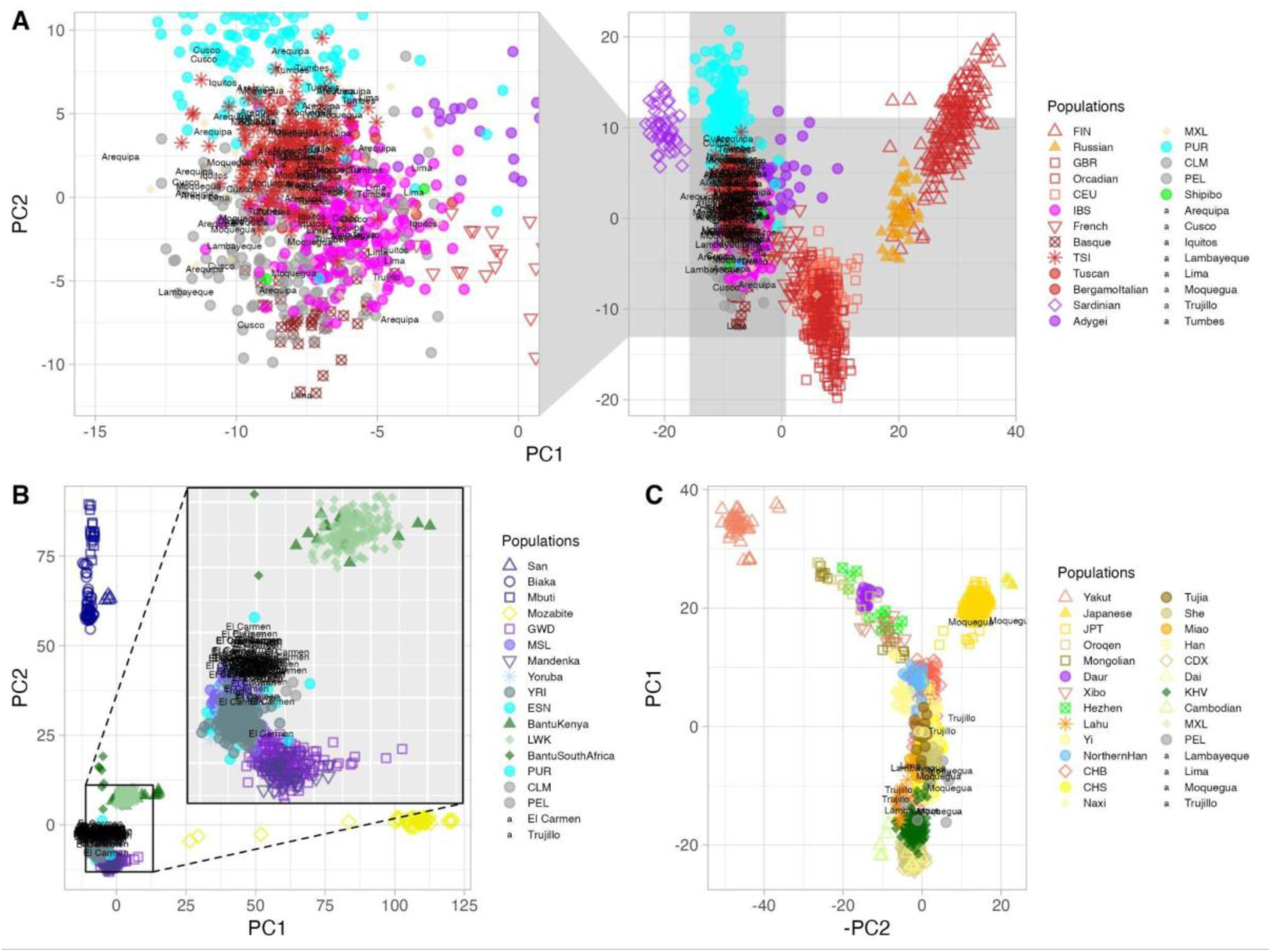
Ancestry Specific Principal Component Analyses (asPCA) to explore subcontinental relationships between Peruvian individuals and worldwide populations. We explored three continental ancestries: A) European, B) African, and C) East Asian. Native American ancestries are shown in Figure 1B.

**Figure S6.**
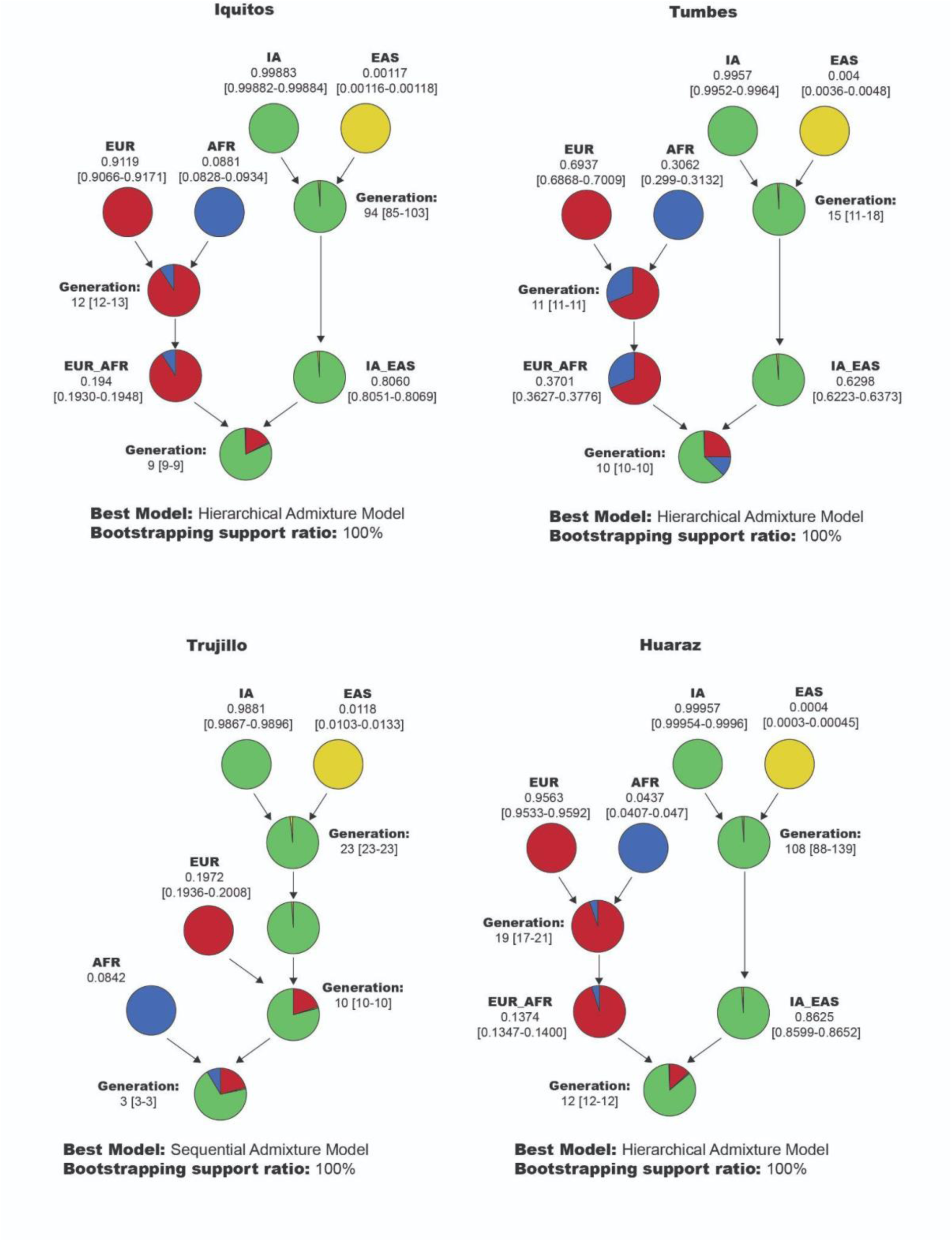
*HiercharchyMix* results for four urban Peruvian areas based on local ancestry results based on four ancestral ancestries. Admixture models inferred for Iquitos, Tumbes, Trujillo, and Huaraz. Each plot represents the admixture model for each population inferred after keeping for ancestry segments greater than 2cM and performing 10000 bootstrap replicates.

**Figure S7.**
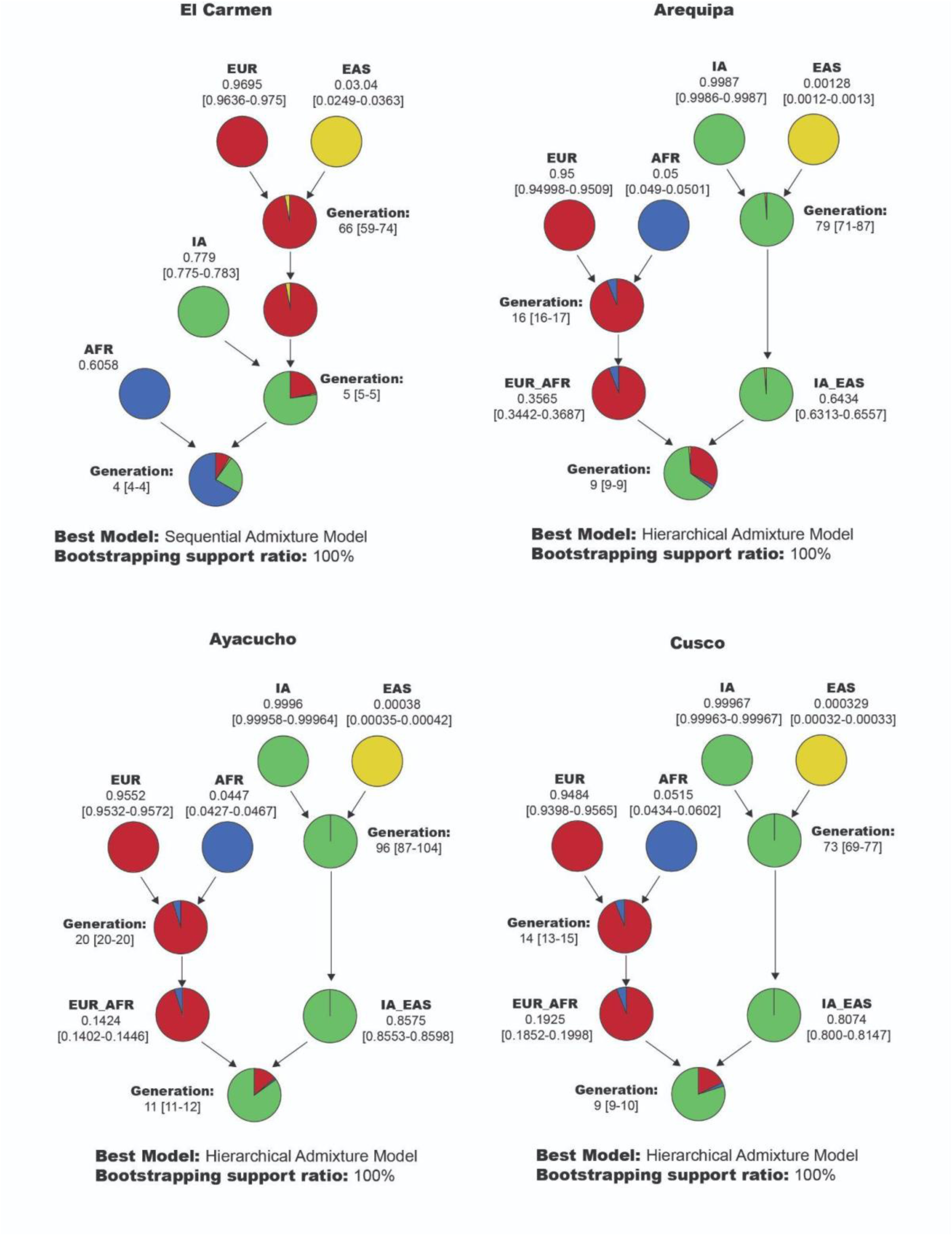
*HiercharchyMix* results for four urban Peruvian areas based on local ancestry results based on four ancestral ancestries. Admixture models inferred for El Carmen, Arequipa, Ayacucho, and Cusco. Each plot represents the admixture model for each population inferred after keeping for ancestry segments greater than 2cM and performing 10000 bootstrap replicates.

**Figure S8.**
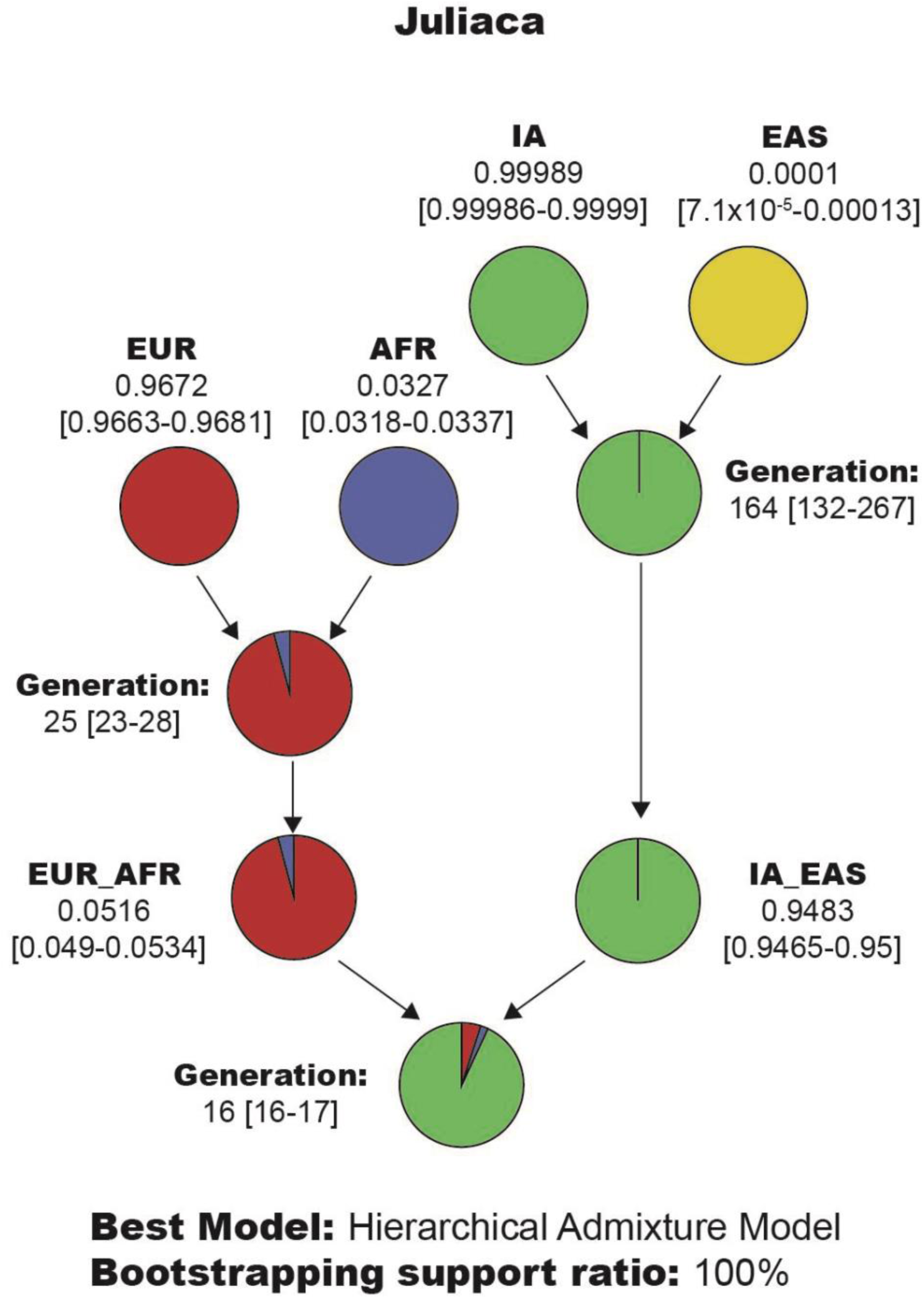
*HiercharchyMix* results for an urban Peruvian area based on local ancestry results based on four ancestral ancestries. Admixture models inferred for Juliaca. Each plot represents the admixture model for each population inferred after keeping for ancestry segments greater than 2cM and performing 10000 bootstrap replicates.

**Figure S9.**
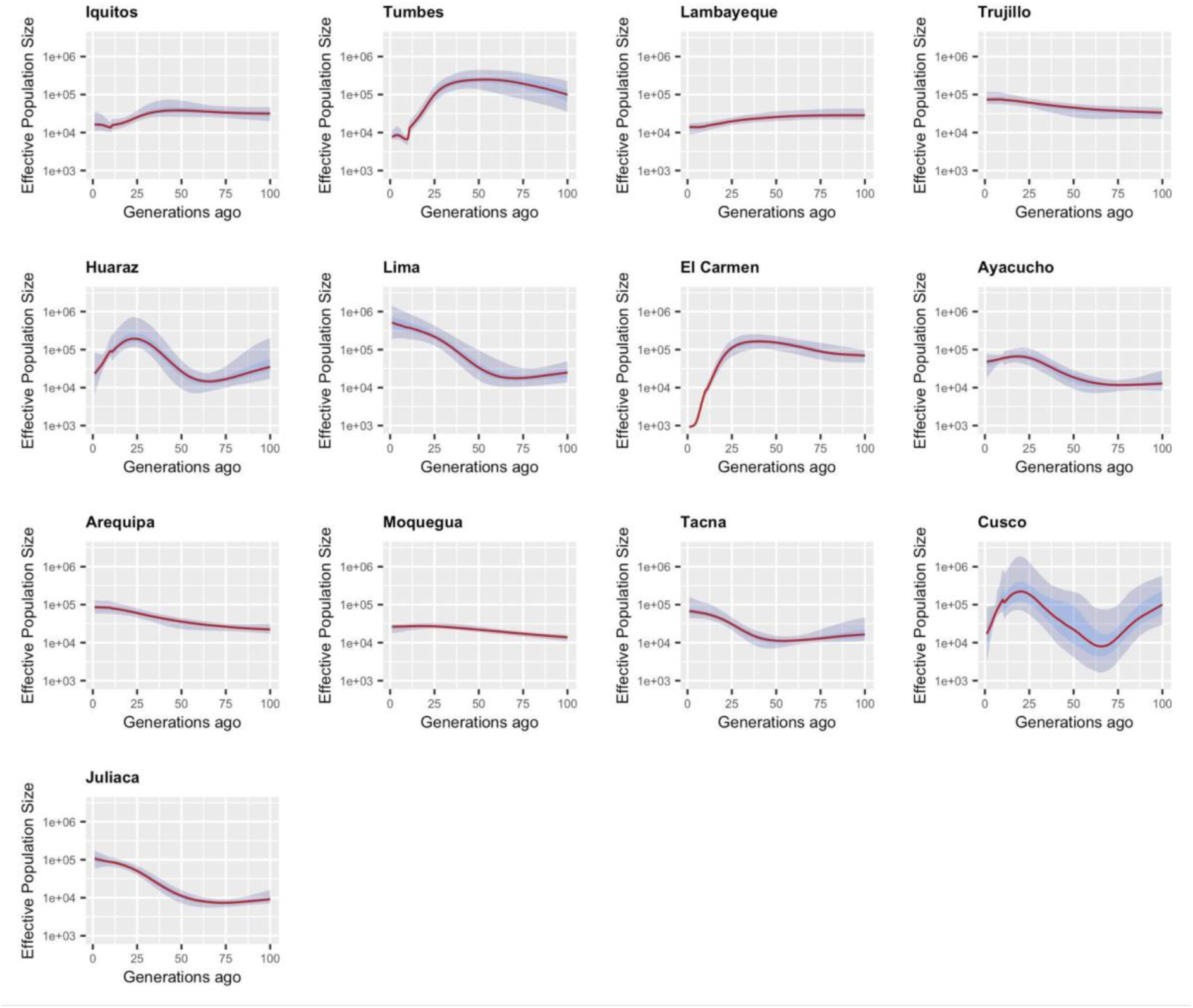
Effective population size trajectory inferred from IBD distribution with HapNe. The red line represents the Maximum likelihood estimate, and shared areas, dark and light blue, represent the 50% and 95% confidence intervals, respectively. The y-axis is in the log scale.

**Figure S10.**
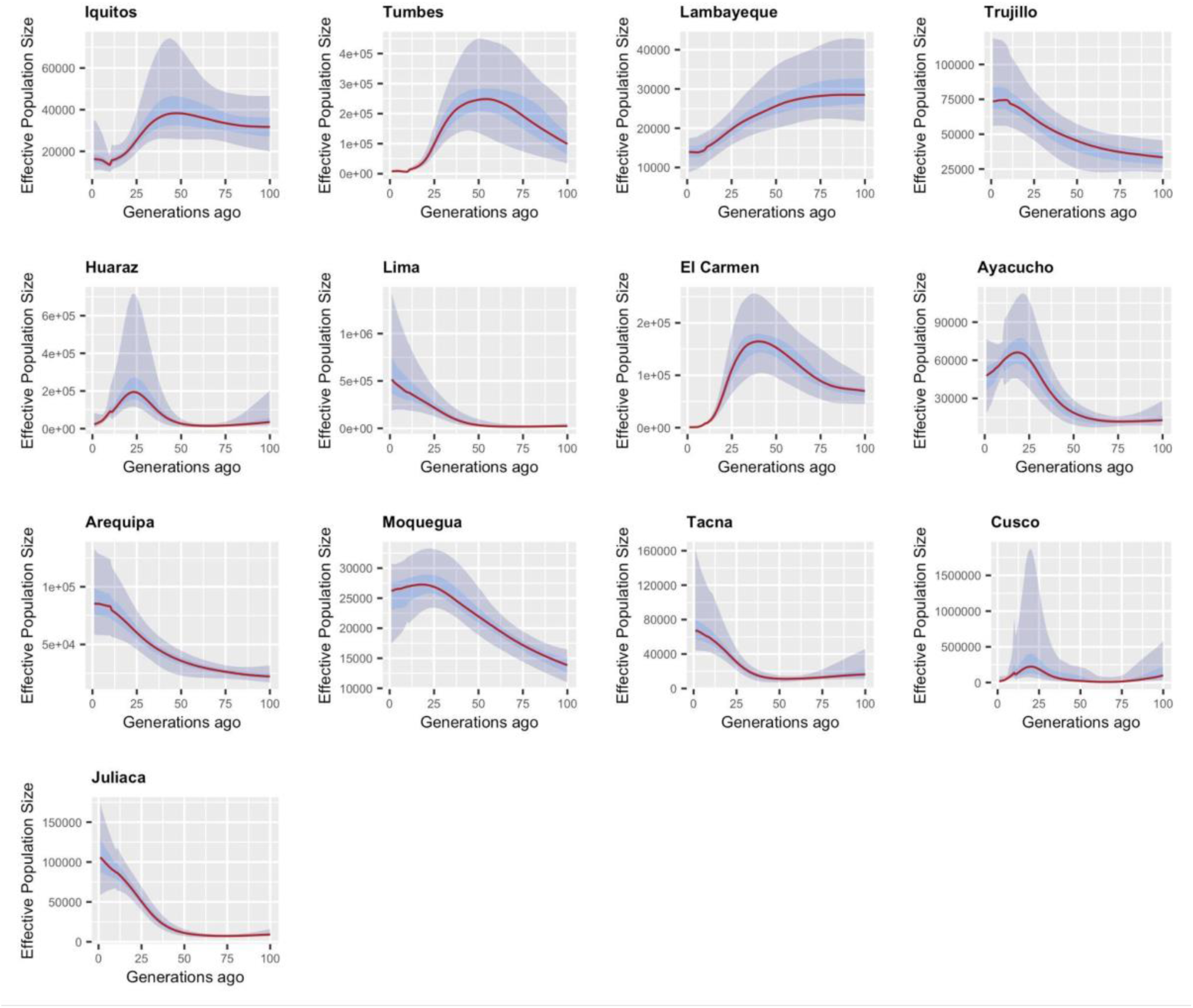
Effective population size trajectory inferred from IBD distribution with HapNe. The red line represents the Maximum likelihood estimate, and shared areas, dark and light blue, represent the 50% and 95% confidence intervals, respectively. The y-axis is on different scales to show the distribution details.

**Figure S11.**
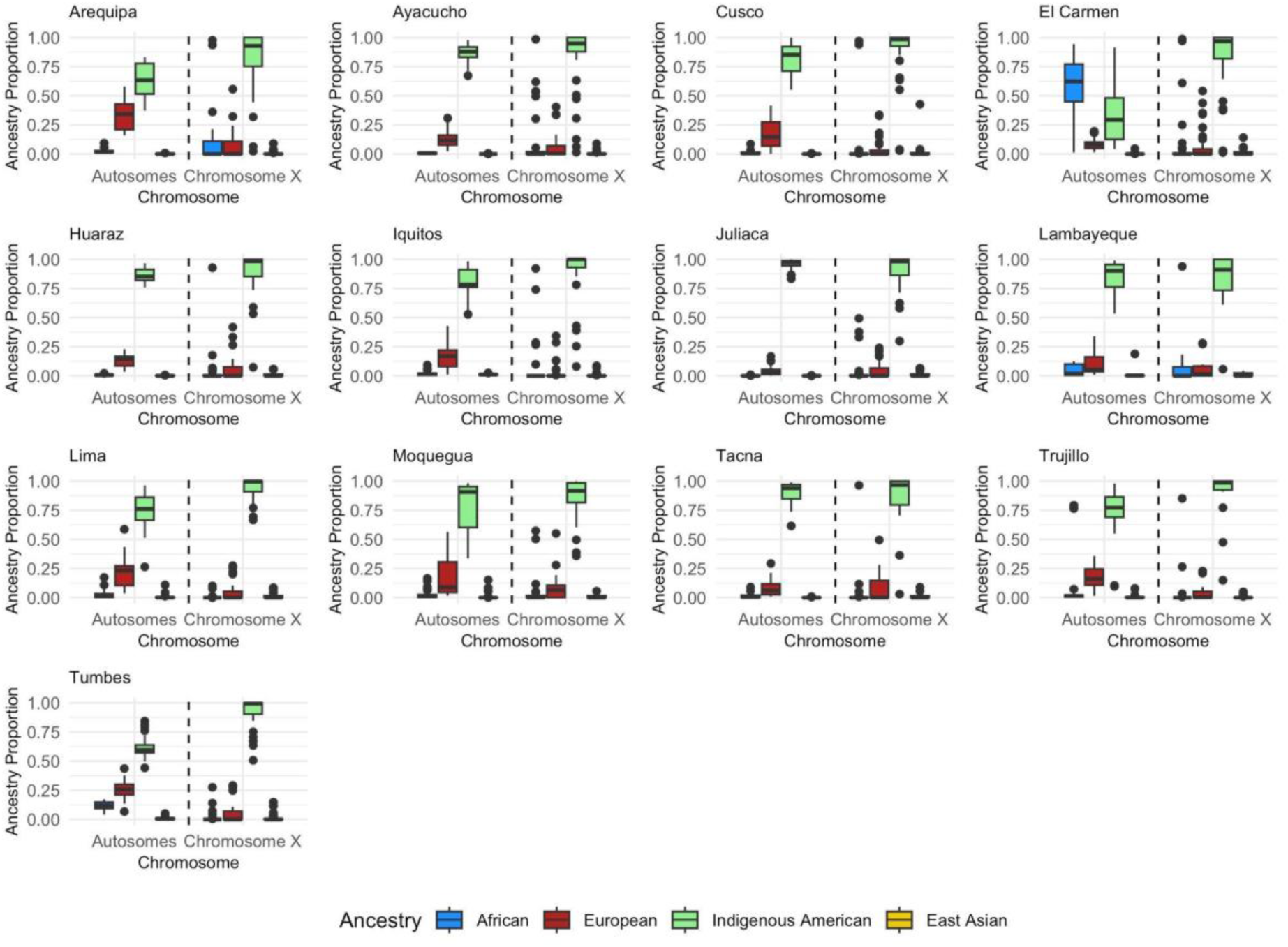
Genome-wide ancestry proportions for chromosome X (gray shade) and autosomes in 13 Admixed Peruvian populations. Ancestry proportions were inferred using ADMIXTURE.

## Notes

### Competing Interest Statement

The authors have declared no competing interest.

## References

1. Mills, M. C. & Rahal, C. The GWAS Diversity Monitor tracks diversity by disease in real time. Nat Genet 52, 242–243 (2020).

2. Chacón-Duque, J.-C. et al. Latin Americans show wide-spread Converso ancestry and imprint of local Native ancestry on physical appearance. Nat Commun 9, 5388 (2018).

3. Borda, V. et al. Genetics of Latin American Diversity (GLAD) Project: insights into population genetics and association studies in recently admixed groups in the Americas. bioRxiv 2023–01 (2023).

4. Homburger, J. R. et al. Genomic Insights into the Ancestry and Demographic History of South America. PLoS Genet 11, e1005602 (2015).

5. Gouveia, M. H. et al. Origins, Admixture Dynamics, and Homogenization of the African Gene Pool in the Americas. Molecular Biology and Evolution 37, 1647–1656 (2020).

6. Ongaro, L. et al. The Genomic Impact of European Colonization of the Americas. Current Biology 29, 3974–3986.e4 (2019).

7. Medina-Muñoz, S. G., et al. Demographic modeling of admixed Latin American populations from whole genomes. The American Journal of Human Genetics 110, 1804–1816 (2023).

8. Harris, D. N. et al. Evolutionary genomic dynamics of Peruvians before, during, and after the Inca Empire. Proc Natl Acad Sci USA 115, E6526–E6535 (2018).

9. Borda, V. et al. The genetic structure and adaptation of Andean highlanders and Amazonians are influenced by the interplay between geography and culture. Proc Natl Acad Sci USA 117, 32557–32565 (2020).

10. Sohail, M. et al. Mexican Biobank advances population and medical genomics of diverse ancestries. Nature 622, 775–783 (2023).

11. The 1000 Genomes Project Consortium et al. A global reference for human genetic variation. Nature 526, 68–74 (2015).

12. Sandoval, J. R. et al. Tracing the genomic ancestry of Peruvians reveals a major legacy of pre-Columbian ancestors. J Hum Genet 58, 627–634 (2013).

13. Iannacone, G. C. et al. Peruvian genetic structure and their impact in the identification of Andean missing persons: A perspective from Ayacucho. Forensic Science International: Genetics Supplement Series 3, e291–e292 (2011).

14. Nakatsuka, N. et al. A Paleogenomic Reconstruction of the Deep Population History of the Andes. Cell 181, 1131–1145.e21 (2020).

15. Urban, M. & Barbieri, C. North and South in the ancient Central Andes: Contextualizing the archaeological record with evidence from linguistics and molecular anthropology. Journal of Anthropological Archaeology 60, 101233 (2020).

16. Cook, N. D. Patrones de migración indígena en el Virreinato del Perú: mitayos, mingas y forasteros. HISTORICA 13, 125–152 (1989).

17. Escobar, G. & Beall, C. M. Contemporary Patterns of Migration in the Central Andes. Mountain Research and Development 2, 63 (1982).

18. Anna, T. E. The Peruvian Declaration of Independence: Freedom by Coercion. Journal of Latin American Studies 7, 221–248 (1975).

19. OʹToole, R. S. Bound Lives: Africans, Indians, and the Making of Race in Colonial Peru. (University of Pittsburgh Press, 2012). doi:10.2307/j.ctt5hjpjn.

20. Gonzales, M. J. Chinese Plantation Workers and Social Conflict in Peru in the Late Nineteenth Century. Journal of Latin American Studies 21, 385–424 (1989).

21. Takenaka, A. The Japanese in Peru: History of Immigration, Settlement, and Racialization. Latin American Perspectives 31, 77–98 (2004).

22. Rodríguez Mega, E. How the mixed-race mestizo myth warped science in Latin America. Nature 600, 374–378 (2021).

23. Menton, M. & Cronkleton, P. Migration and Forests in the Peruvian Amazon: A Review. (Center for International Forestry Research (CIFOR), 2019). doi:10.17528/cifor/007305.

24. INEI. Perú: Perfil Sociodemográfico Informe Nacional. (2018).

25. Contreras, C. & Cueto, M. HISTORIA DEL PERÚ CONTEMPORÁNEO. (Instituto de Estudios Peruanos, Peru, 2024).

26. Guio, H. et al. The Peruvian Genome Project: expanding the global pool of genome diversity from South America. Preprint at 10.1101/2024.05.05.24306840 (2024).

27. Byrska-Bishop, M. et al. High-coverage whole-genome sequencing of the expanded 1000 Genomes Project cohort including 602 trios. Cell 185, 3426–3440.e19 (2022).

28. Bergström, A. et al. Insights into human genetic variation and population history from 929 diverse genomes. Science 367, eaay5012 (2020).

29. Alexander, D. H., Novembre, J. & Lange, K. Fast model-based estimation of ancestry in unrelated individuals. Genome Research 19, 1655–1664 (2009).

30. Moreno-Estrada, A. et al. Reconstructing the Population Genetic History of the Caribbean. PLoS Genet 9, e1003925 (2013).

31. Browning, S. R. et al. Local Ancestry Inference in a Large US-Based Hispanic/Latino Study: Hispanic Community Health Study/Study of Latinos (HCHS/SOL). G3 Genes|Genomes|Genetics 6, 1525–1534 (2016).

32. Lawson, D. J., Hellenthal, G., Myers, S. & Falush, D. Inference of Population Structure using Dense Haplotype Data. PLoS Genet 8, e1002453 (2012).

33. Bonfiglio, G. Introducción al estudio de la inmigración europea en el Perú. Apuntes: Revista de Ciencias Sociales 93–127 (2015).

34. Bowser, F. P. The African Slave in Colonial Peru, 1524-1650. (Stanford University Press, Stanford, Calif, 1974).

35. Sobrevilla Perea, N. The Abolition of Slavery in the South American Republics. Slavery & Abolition 44, 90–108 (2023).

36. Hellenthal, G. et al. A Genetic Atlas of Human Admixture History. Science 343, 747– 751 (2014).

37. Zhang, S. et al. Reconstructing complex admixture history using a hierarchical model. Briefings in Bioinformatics 25, bbad540 (2024).

38. Wang, R. J., Al-Saffar, S. I., Rogers, J. & Hahn, M. W. Human generation times across the past 250,000 years. Sci. Adv. 9, eabm7047 (2023).

39. Baharian, S. et al. The Great Migration and African-American Genomic Diversity. PLoS Genet 12, e1006059 (2016).

40. Fournier, R., Tsangalidou, Z., Reich, D. & Palamara, P. F. Haplotype-based inference of recent effective population size in modern and ancient DNA samples. Nat Commun 14, 7945 (2023).

41. Ongaro, L. et al. Evaluating the Impact of Sex-Biased Genetic Admixture in the Americas through the Analysis of Haplotype Data. Genes 12, 1580 (2021).

42. Pena, S. D. J., Santos, F. R. & Tarazona-Santos, E. Genetic admixture in Brazil. American J of Med Genetics Pt C 184, 928–938 (2020).

43. Nagar, S. D. et al. Genetic ancestry and ethnic identity in Ecuador. Human Genetics and Genomics Advances 2, 100050 (2021).

44. Kehdy, F. S. G. et al. Origin and dynamics of admixture in Brazilians and its effect on the pattern of deleterious mutations. Proc Natl Acad Sci USA 112, 8696–8701 (2015).

45. Luisi, P. et al. Fine-scale genomic analyses of admixed individuals reveal unrecognized genetic ancestry components in Argentina. PLoS ONE 15, e0233808 (2020).

46. Eyheramendy, S., Martinez, F. I., Manevy, F., Vial, C. & Repetto, G. M. Genetic structure characterization of Chileans reflects historical immigration patterns. Nat Commun 6, 6472 (2015).

47. Adhikari, K., Mendoza-Revilla, J., Chacón-Duque, J. C., Fuentes-Guajardo, M. & Ruiz-Linares, A. Admixture in Latin America. Current Opinion in Genetics & Development 41, 106–114 (2016).

48. Alvim, I. et al. The need to diversify genomic studies: Insights from Andean highlanders and Amazonians. Cell 187, 4819–4823 (2024).

49. The Oxford Handbook of Latin American History. (Oxford University Press, Oxford; New York, 2011).

50. Aguirre, C. Breve Historia de La Esclavitud En El Perú: Una Herida Que No Deja de Sangrar. (Fondo Editorial del Congreso del Perú, Lima, 2005).

51. Golash-Boza, T. M. Yo Soy Negro: Blackness in Peru. (University Press of Florida, Gainesville, 2012).

52. Instituto Nacional de Estadistica e Informatica. Situacion de la Poblacon Peruana 2024: Una mirada de la diversidad étnica. (2024).

53. Pedersen, D., Tremblay, J., Errázuriz, C. & Gamarra, J. The sequelae of political violence: Assessing trauma, suffering and dislocation in the Peruvian highlands. Social Science & Medicine 67, 205–217 (2008).

54. Loret De Mola, C., et al. The effect of rural-to-urban migration on social capital and common mental disorders: PERU MIGRANT study. Soc Psychiatry Psychiatr Epidemiol 47, 967–973 (2012).

55. Falaris, E. M. The Determinants of Internal Migration in Peru: An Economic Analysis. Economic Development and Cultural Change 27, 327–341 (1979).

56. Vicuña, L. et al. Novel loci and Mapuche genetic ancestry are associated with pubertal growth traits in Chilean boys. Hum Genet 140, 1651–1661 (2021).

57. Purcell, S. et al. PLINK: A Tool Set for Whole-Genome Association and Population-Based Linkage Analyses. The American Journal of Human Genetics 81, 559–575 (2007).

58. Manichaikul, A. et al. Robust relationship inference in genome-wide association studies. Bioinformatics 26, 2867–2873 (2010).

59. Zheng, X. et al. A high-performance computing toolset for relatedness and principal component analysis of SNP data. Bioinformatics 28, 3326–3328 (2012).

60. Leal, T. P. et al. NAToRA, a relatedness-pruning method to minimize the loss of dataset size in genetic and omics analyses. Computational and Structural Biotechnology Journal 20, 1821–1828 (2022).

61. Delaneau, O., Zagury, J.-F., Robinson, M. R., Marchini, J. L. & Dermitzakis, E. T. Accurate, scalable and integrative haplotype estimation. Nat Commun 10, 5436 (2019).

62. Conomos, M. P., Miller, M. B. & Thornton, T. A. Robust Inference of Population Structure for Ancestry Prediction and Correction of Stratification in the Presence of Relatedness. Genet. Epidemiol. 39, 276–293 (2015).

63. Weir, B. S. & Cockerham, C. C. Estimating F-Statistics for the Analysis of Population Structure. Evolution 38, 1358 (1984).

64. Zhou, Y., Browning, S. R. & Browning, B. L. A Fast and Simple Method for Detecting Identity-by-Descent Segments in Large-Scale Data. The American Journal of Human Genetics 106, 426–437 (2020).

65. Hilmarsson, H., et al. High Resolution Ancestry Deconvolution for Next Generation Genomic Data. http://biorxiv.org/lookup/doi/10.1101/2021.09.19.460980 (2021) doi:10.1101/2021.09.19.460980.

